# Click-Prep: An Interactive Data Preparation Tool for Click-qPCR

**DOI:** 10.64898/2026.08.20.745930

**Authors:** Azusa Kubota, Atsushi Tajima

## Abstract

Click-qPCR is a browser-based application for relative qPCR analysis that requires a tidy-format CSV file containing four columns: sample, group, gene, and Cq. Preparing this input from qPCR instrument output typically requires manual reformatting and calculation of mean Cq values for technical replicates. To simplify this process, we developed Click-Prep (https://kubo-azu.shinyapps.io/Click-Prep/), an interactive web-based application designed specifically to create Click-qPCR input files. Click-Prep imports CSV, TXT, TSV, and XLS/XLSX files and supports skipping of instrument-generated metadata rows, interactive column mapping, and manual assignment of experimental groups. Users can review technical-replicate measurements, exclude selected rows according to predefined quality-control criteria, and calculate mean Cq values for each sample– group–target combination. Missing or nonnumeric Cq values are flagged for review and must be resolved before the mean is calculated. Click-Prep can also combine compatible formatted CSV files, such as datasets obtained from separate qPCR plates. The resulting dataset is exported as a standardized CSV file containing the four fields required by Click-qPCR. By integrating these operations into a guided browser-based workflow, Click-Prep enables users to prepare Click-qPCR input files rapidly and consistently without programming.

**Key features:**

- Creates standardized Click-qPCR input files containing sample, group, gene, and Cq columns.
- Imports tabular qPCR instrument output and supports metadata-row skipping, interactive column mapping, and manual group assignment.
- Supports user-controlled review and exclusion of technical-replicate measurements and calculation of mean Cq values.
- Combines compatible mean Cq datasets from multiple qPCR plates into a single Click-qPCR input file.

**Graphical overview:** 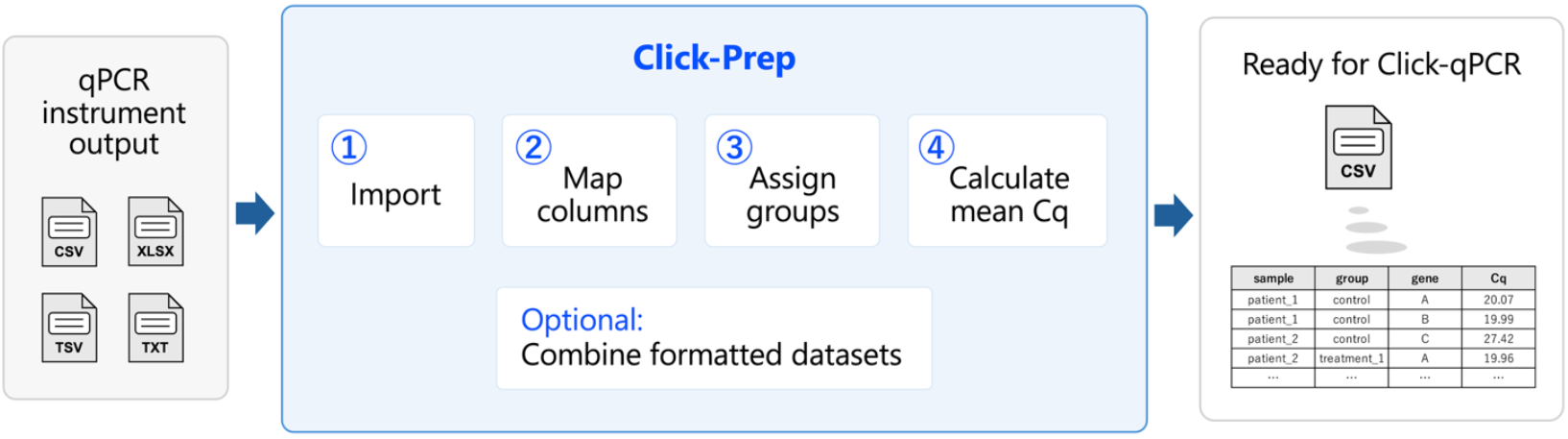

## Background

Real-time quantitative PCR (qPCR) is widely used for the relative quantification of gene expression and DNA copy number variations (CNVs) [1]. In both applications, target Cq values can be normalized against appropriate reference genes or genomic loci, and relative differences between experimental groups can be evaluated using the ΔCq or ΔΔCq method [2,3]. Reliable analysis therefore depends not only on experimental design and amplification quality but also on accurate and transparent processing of the Cq values generated by qPCR instruments. The MIQE 2.0 guidelines emphasize that data-processing procedures, including the handling of technical replicates, missing values, and excluded measurements, should be transparently documented to support the reliability and reproducibility of qPCR experiments [4].

We previously developed Click-qPCR, a browser-based application that enables users to perform ΔCq and ΔΔCq calculations, statistical comparisons, and interactive visualization without programming [5]. Click-qPCR requires a tidy-format CSV file containing four variables: sample, group, gene, and Cq. However, tabular files exported from qPCR instruments do not necessarily conform to this format and may contain instrument-specific metadata, different column names, or technical replicate measurements. Moreover, experimental-group information may be absent from the exported file. Users must therefore reorganize the data, assign group labels, review technical replicates, calculate mean Cq values, and, when necessary, combine data obtained from multiple plates before analysis. Performing these operations manually in spreadsheet software can be time-consuming and may introduce transcription, labeling, or formatting errors.

To facilitate the rapid and consistent preparation of Click-qPCR input files, we developed Click-Prep, an interactive web-based application that converts tabular qPCR instrument output into the format required by Click-qPCR. Click-Prep supports tabular input in CSV, TXT, TSV, and XLS/XLSX formats; skipping of leading instrument-metadata rows; interactive column mapping; and manual assignment of experimental groups. Users can also visually inspect technical replicate measurements, manually exclude selected rows, calculate mean Cq values for each sample and target, and combine multiple formatted CSV files. By making these preparatory operations explicit within a standardized workflow, Click-Prep supports transparent and consistent handling of qPCR data before downstream analysis. Together, Click-Prep and Click-qPCR provide a continuous browser-based workflow from data preparation to statistical analysis and visualization.

**Table 1.** Required packages for Click-Prep application.

| Type | Software/dataset/resource | Version | Date | License | Access (free or paid) |
| --- | --- | --- | --- | --- | --- |
| Software | R [6] | 4.6.1 | 2026.06.24 | GNU GPL | Free |
| Software | dplyr [7] | 1.2.1 | 2026.04.03 | MIT | Free |
| Software | DT [8] | 0.34.0 | 2025.09.02 | MIT | Free |
| Software | readr [9] | 2.2.0 | 2026.02.19 | MIT | Free |
| Software | readxl [10] | 1.5.0 | 2026.05.16 | MIT | Free |
| Software | shiny [11] | 1.14.0 | 2026.06.21 | MIT | Free |
| Software | shinyjs [12] | 2.1.1 | 2026.01.15 | MIT | Free |

## Software and datasets

The Click-Prep application was developed using the R programming language and Shiny framework to provide an interactive web interface. The required packages for Click-Prep are shown in Table 1. The application’s source code is publicly available at https://github.com/kubo-azu/Click-Prep.

## Procedure

### A. Running the application

#### Option 1: Run the hosted application in a web browser

Open https://kubo-azu.shinyapps.io/Click-Prep/ in a standard web browser.

#### Option 2: Run directly from GitHub

Click-Prep can be launched directly from GitHub using shiny::runGitHub():

~~~
```R
if (!requireNamespace(“shiny”, quietly = TRUE)) {
    install.packages(“shiny”)
}
shiny::runGitHub(“kubo-azu/Click-Prep”)
```
~~~

Required packages that are not already installed may need to be installed before the application can start; see Table 1 for details. Because shiny::runGitHub() does not restore the package versions recorded in renv.lock, cloning the repository and using renv::restore () is required for reproducible local execution.

#### Option 3: Clone and run the application locally

1. Clone the repository: ```Terminal git clone https://github.com/kubo-azu/Click-Prep.git ```
2. Open the cloned directory in RStudio by opening Click-Prep.Rproj, or set the cloned directory as the working directory in R.
3. Restore the package environment recorded in renv.lock: ```R if (!requireNamespace(“renv”, quietly = TRUE)) { install.packages(“renv”) } renv::restore() ```
4. Launch the application: ```R shiny::runApp() ```

**Note:** Local execution may be preferable when uploaded data are confidential, regulated, or subject to institutional data-handling requirements.

### B. Importing and formatting qPCR instrument output

1. Upload the instrument output:
  a. Navigate to the “Import and Column Mapping” tab and click the “Browse…” button to upload a CSV, TXT, TSV, or XLS/XLSX file (Figure 1A).
  b. Inspect the “Original Data Preview” table (Figure 1B).
  c. If instrument-generated metadata appears above the column headers, adjust the “Skip first N rows (metadata)” number until the correct header row and data are displayed (Figure 1C). **Critical:** CSV files must be comma-delimited, whereas TXT and TSV files must be tab-delimited. Click-Prep imports only the first worksheet of an XLS/XLSX file.
2. Map the required columns:
  a. In the “Column Mapping” area, select the source columns corresponding to sample names, experimental groups, target genes or genomic loci, and Cq values (Figure 1D).
  b. Confirm that the “Formatted Data Preview” table contains the columns sample, group, gene, and Cq (Figure 1E). **Critical:** Missing or nonnumeric Cq values are displayed as NA and flagged for review. They must be resolved before mean Cq calculation.
3. Manual assignment of experimental groups: Use this option when the instrument output does not contain group information.
  a. Select [Assign groups manually] for the experimental-group column in Step 2-a (Figure 1D). The “Assign Groups Manually” area will appear (Figure 1F).
  b. In Step 1: Define your groups, enter each group name and click the “Add Group” button (Figure 2A).
  c. In Step 2: Assign groups to samples, select one or more samples, choose a group from the “Assign to:” menu, and click the “Apply” button (Figure 2D). Repeat until no samples remain Unassigned.
  d. Confirm the assignments in the “Formatted Data Preview” table (Figure 2E). **Note:** The search field (Figure 2B) and “Select All” button (Figure 2C) can be used to select multiple samples efficiently. The “Clear” button (Figure 2C) erases the assigned group information from the selected samples. **Critical:** Manual group assignment requires source columns corresponding to sample, gene, and Cq. Use consistent group names across samples and plates because capitalization, spacing, and spelling are not standardized automatically.
4. Complete the formatting step: Confirm that “Mapping Complete!” is displayed (Figure 2F). If Cq warnings remain, proceed to the “Manual Data Review and Mean Cq Calculation” tab to exclude the affected measurements or restart with a corrected source file. If required, click the “Download Formatted CSV (All Replicates)” button (Figure 2F) to export the standardized replicate-level dataset.

**Figure 1.**
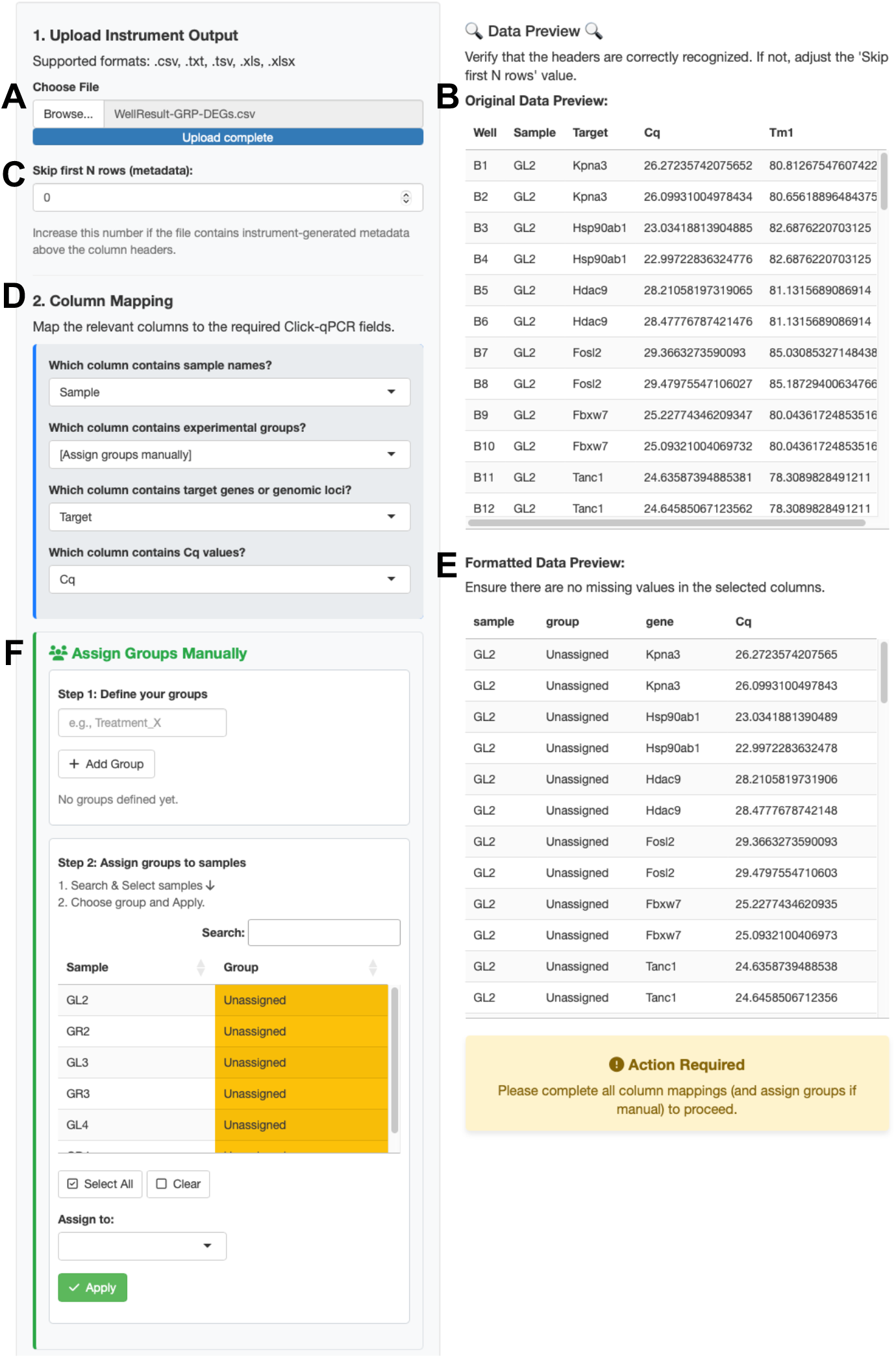
Importing and mapping qPCR instrument output in Click-Prep. **(A)** File upload. **(B)** Preview of the original instrument output. **(C)** Setting the number of leading metadata rows to skip. **(D)** Mapping of source columns to the required Click-qPCR fields. **(E)** Preview of the formatted data. **(F)** Manual group assignment when group information is absent from the source file.

**Figure 2.**
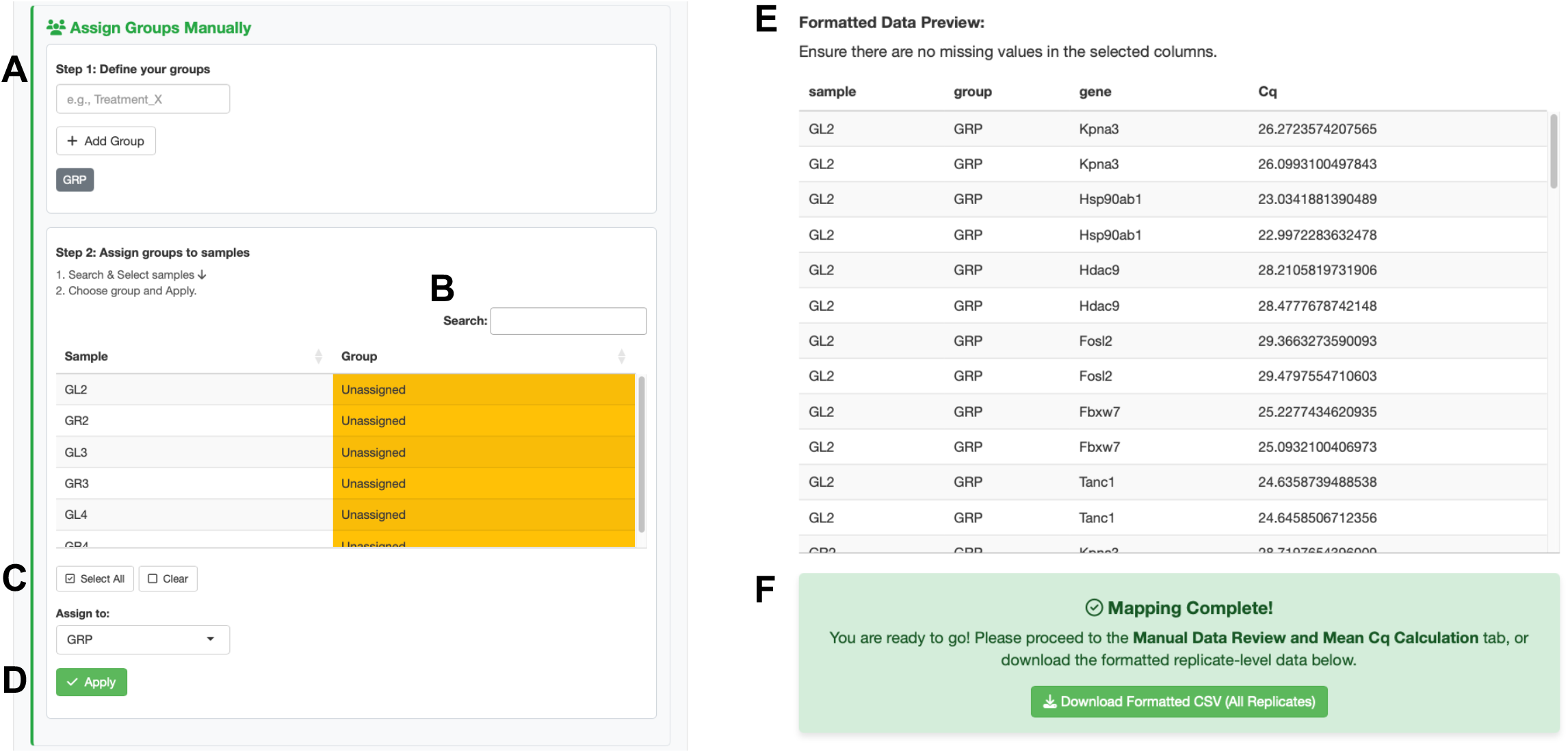
Manual assignment of experimental groups. **(A)** Definition of group names. **(B)** Sample search. **(C)** Sample-selection controls. **(D)** Assignment of selected samples to a group. **(E)** Preview after group assignment. **(F)** Completion message and download of the formatted replicate-level dataset.

### C. Reviewing replicate measurements and calculating mean Cq values

1. Review and exclude replicate measurements:
  a. Navigate to the “Manual Data Review and Mean Cq Calculation” tab and inspect the replicate-level data (Figure 3A). Missing or nonnumeric Cq values are displayed as NA with a warning.
  b. If a measurement should be excluded, select the corresponding row and click the “Exclude Selected Rows” button. Repeat this step as necessary.
  c. If required, click the “Download Retained Replicate Data” button to save the replicate-level dataset after exclusion. **Critical:** Click-Prep does not automatically identify outliers. Measurements should be excluded only according to predefined quality-control criteria. All missing or invalid Cq values must be resolved before mean calculation.
2. Calculate and export mean Cq values:
  a. Click the “Calculate Mean Cq” button (Figure 3A). If invalid Cq values remain, Click-Prep displays an error and does not perform the calculation.
  b. Confirm the results in the “Mean Cq Data Preview” table (Figure 3B).
  c. Click the “Download Mean Cq Dataset (Ready for Click-qPCR)” button to export the resulting CSV file. Click-Prep calculates the arithmetic mean of the retained measurements for each sample–group–gene combination and exports the columns sample, group, gene, and Cq.

**Figure 3.**
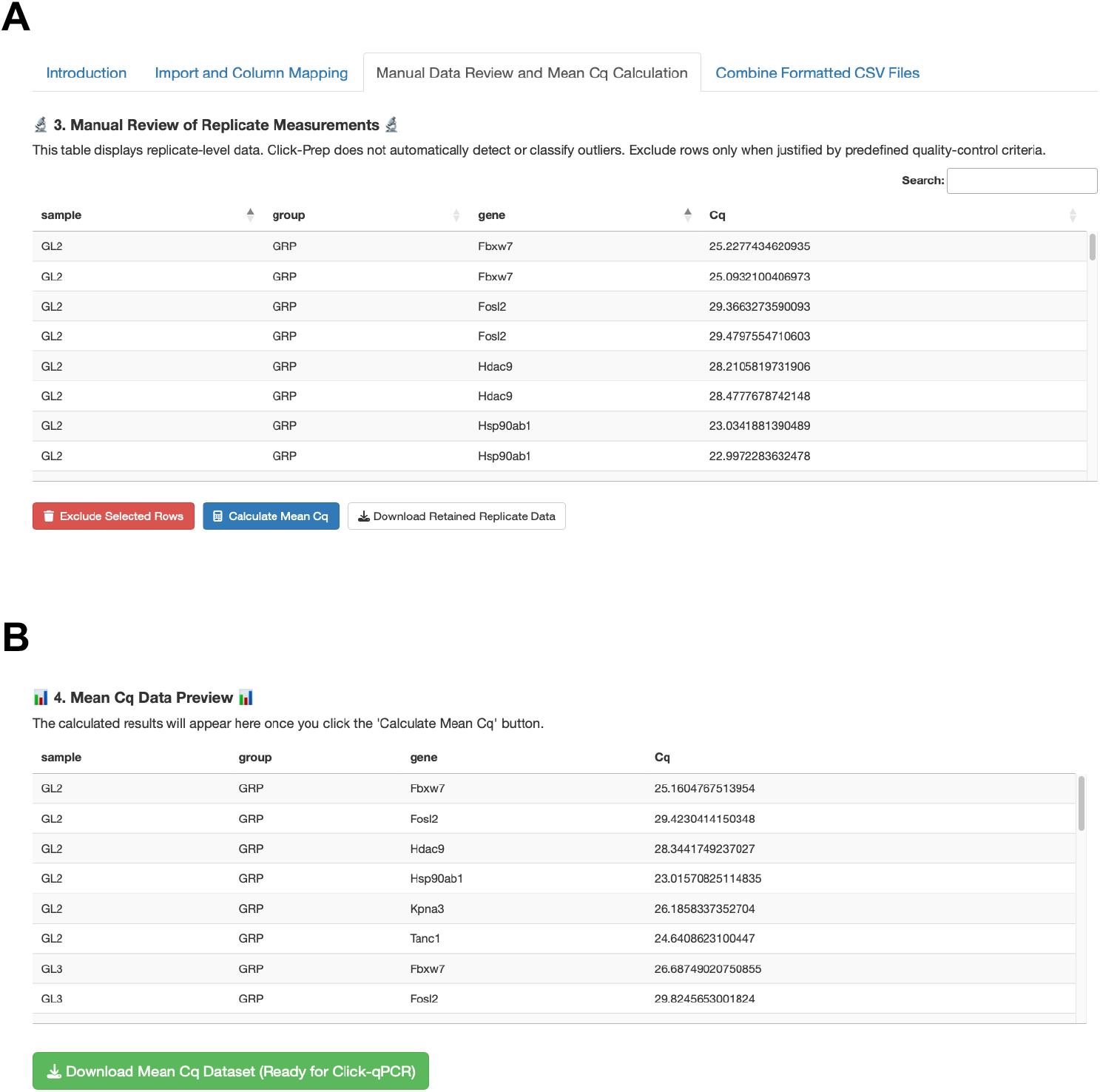
Review of technical replicates and calculation of mean Cq values. **(A)** Review and optional exclusion of replicate-level measurements and calculation of mean Cq values. **(B)** Preview and download of the mean Cq dataset calculated from the retained measurements.

### D. Combining formatted CSV files

Use this optional function to combine mean Cq datasets from separate qPCR plates.

1. Upload and review formatted CSV files:
  a. Navigate to the “Combine Formatted CSV Files” tab and click the “Browse…” button (Figure 4A) to upload one or more formatted CSV files. Additional files can be appended by repeating this step.
  b. Confirm the uploaded filenames and inspect the combined dataset in the “Merged Data Preview” table (Figure 4B). Click-Prep accepts files containing exactly the columns sample, group, gene, and Cq, in this order. The required fields must not contain missing values, and all Cq values must be numeric. If any file fails these checks, an error message is displayed, and the combined dataset is not generated.
2. Export or reset the combined dataset:
  a. After reviewing the preview, click the “Download Merged CSV” button (Figure 4C) to export the combined dataset.
  b. To remove all uploaded files and begin a new combination, click the “Reset Files” button. The exported mean Cq dataset can be used directly as input for Click-qPCR.

**Figure 4.**
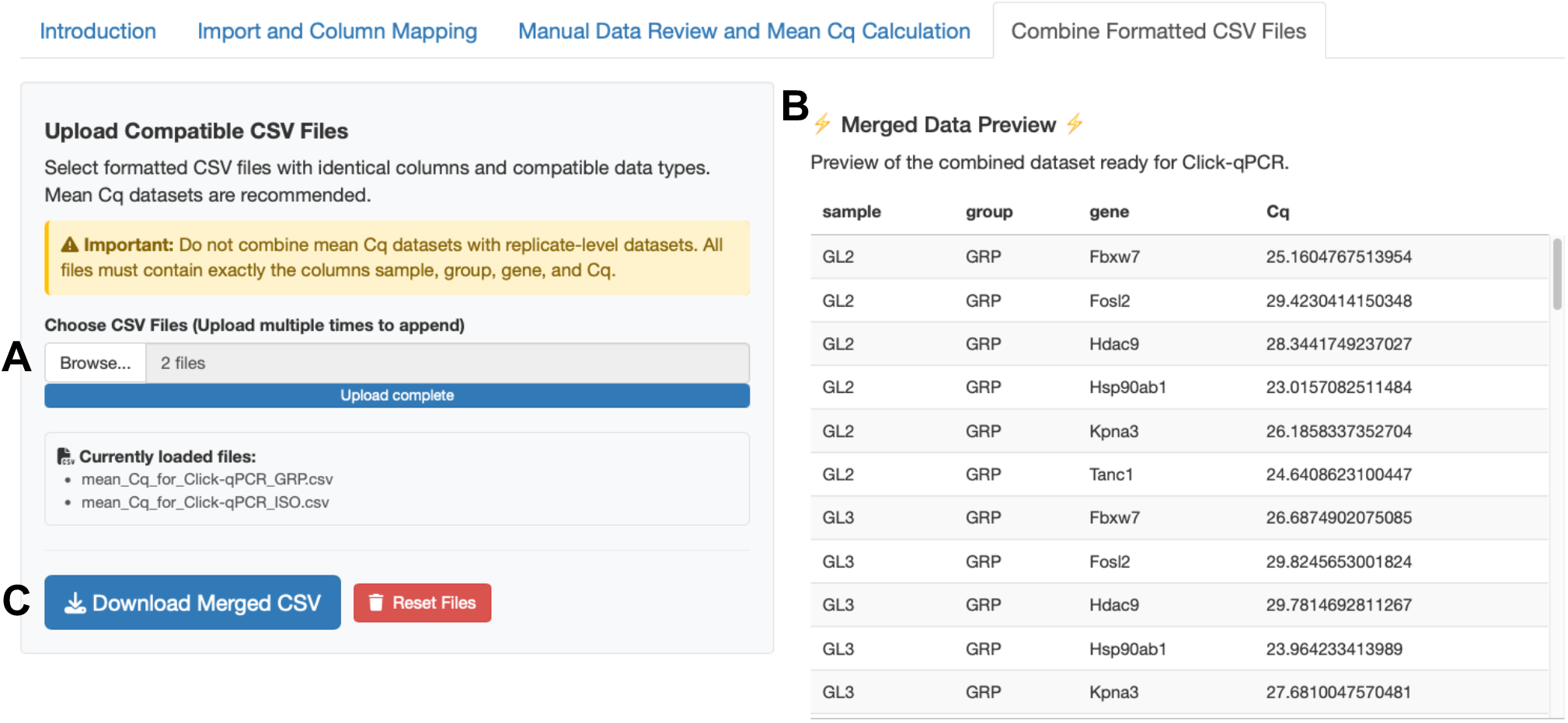
Combining formatted CSV files. **(A)** Upload and management of compatible CSV files. **(B)** Preview of the combined dataset. **(C)** Download of the merged CSV file or reset of the uploaded files.

## General notes and troubleshooting

### General notes

1. Click-Prep processes tabular qPCR output containing Cq values. It does not assess raw fluorescence data, amplification or melting curves, amplification efficiency, assay specificity, controls, or reference-target suitability. These aspects must be evaluated separately using appropriate qPCR quality-control procedures.
2. Click-Prep does not directly import proprietary qPCR experiment files, RDML/XML files, or report files. Results must first be exported as a comma-delimited CSV, tab-delimited TXT/TSV, or XLS/XLSX file containing sample identifiers, target names, and Cq values.
3. Click-Prep does not automatically identify outliers. Missing or nonnumeric Cq values are flagged for review, but measurement exclusion remains the user’s responsibility and should follow predefined quality-control criteria.
4. Excluded rows are removed from the working dataset, but Click-Prep does not record the excluded measurements or reasons for exclusion. Users should retain the original data and document all exclusions and the criteria applied.
5. Mean Cq values are calculated as the arithmetic mean of the retained technical replicates for each sample– group–gene combination. Click-Prep does not assess replicate variability or adequacy, and technical replicates must not be treated as independent biological replicates.
6. The CSV-combination function concatenates compatible datasets by rows. It does not detect or correct plate effects, batch effects, duplicate sample identifiers, or inconsistent annotations. Users should ensure that datasets are consistent and scientifically appropriate to combine.
7. Uploaded data are processed within the active application session. Click-Prep does not intentionally write uploaded qPCR data to persistent application storage, and temporary files created by the application are removed when they are no longer required or when the session ends.
8. Click-Prep facilitates standardized preparation of Click-qPCR input files but does not by itself ensure compliance with the MIQE 2.0 guidelines [4]. Experimental design, assay validation, quality control, normalization, statistical analysis, and reporting remain the user’s responsibility.

### Troubleshooting

Problem 1: The column headers are not recognized correctly after the file is uploaded.

Possible cause: The instrument output contains metadata rows above the actual column headers, or the file delimiter is not supported.

Solution: In the Import and Column Mapping tab, adjust Skip first N rows (metadata) until the actual header row is correctly displayed in the Original Data Preview. CSV files must be comma-delimited, whereas TXT and TSV files must be tab-delimited. If necessary, open the file in spreadsheet software and export it in one of these formats before uploading it again.

Problem 2: Mean Cq values cannot be calculated.

Possible cause: One or more rows contain missing, nonnumeric, or nonfinite Cq values. Such entries may originate from values such as blank cells, NA, N/A, or Undetermined in the instrument output.

Solution: Review the warning and the replicate-level data in the Manual Data Review and Mean Cq Calculation tab. Exclude the affected rows if exclusion is justified, or click Start Over / Reset All and upload a corrected input file. Click-Prep does not calculate mean Cq values while invalid Cq values remain.

Problem 3: Formatted CSV files cannot be combined.

Possible cause: The uploaded files do not have identical structures, contain missing or nonnumeric values, or include a mixture of replicate-level and mean Cq datasets.

Solution: Confirm that every file contains exactly the columns sample, group, gene, and Cq, in this order. Verify that all Cq values are numeric and that the other required fields are complete. Do not combine replicate-level datasets with mean Cq datasets. Mean Cq files generated by Click-Prep are recommended for this function.

Problem 4: The locally installed application fails to start, or renv reports that the project library and lockfile are out of sync.

Possible cause: The active R version or installed package versions differ from those used for Click-Prep version 1.2.1.

Solution: Confirm that R 4.6.1 is active using R.version.string in the R console. If multiple R versions are installed, select R 4.6.1 using RStudio or a version manager such as rig:

```Terminal
rig rstudio 4.6 Click-Prep.Rproj
```

Then, restore and check the project environment:

```R

renv::restore()

renv::status()

```

If renv itself is inconsistent, run renv::restore(packages = “renv”), restart R, and repeat renv::restore().

## Acknowledgements

Writing – original draft: A.K. Writing – review & editing: A.T. Conceptualization: A.K. Software: A.K. Validation: A.K., A.T. Funding acquisition: A.K., A.T. Supervision: A.T.

This study was supported by JST SPRING (grant number: JPMJSP2135). We thank the Center for Biomedical Research and Education of Kanazawa University for their technical assistance.

## Competing interests

The authors declare that they have no competing interests.

## Notes

### Competing Interest Statement

The authors have declared no competing interest.

